# Differential Akt Signaling Induced by Tumstatin and Endostatin in Human Endothelial Cells

**DOI:** 10.64898/2026.06.09.731223

**Authors:** Valerie S. Kalluri, Raghu Kalluri

## Abstract

Tumstatin and Endostatin are endogenous extracellular matrix-derived inhibitors of angiogenesis generated from the non-collagenous domains of type IV and type XVIII collagens, respectively. Although both molecules suppress angiogenesis and tumor growth in vivo, previous studies demonstrated that they engage distinct endothelial integrin receptors and activate different intracellular signaling pathways. In particular, Tumstatin inhibits endothelial proliferation through suppression of the focal adhesion kinase (FAK)/phosphatidylinositol 3-kinase (PI3K)/Akt/mTOR pathway, whereas Endostatin primarily inhibits endothelial migration through α5β1 integrin-dependent signaling. Here we evaluated a key mechanistic distinction between these two angiogenesis inhibitors by examining Akt phosphorylation in human umbilical vein endothelial cells (HUVEC) cultured on fibronectin. Validating previous reports, recombinant human Tumstatin reduced Akt phosphorylation whereas recombinant human Endostatin did not alter Akt activation. These findings confirm a defining feature of Tumstatin signaling and reinforce the concept that collagen-derived angiogenesis inhibitors regulate endothelial cell behavior through distinct integrin-dependent mechanisms.

## Introduction

Angiogenesis is a tightly regulated process controlled by a balance between endogenous stimulators and inhibitors of vascular growth^1-4^. Among the most extensively studied endogenous inhibitors are Endostatin, derived from the C-terminal non-collagenous domain of collagen XVIII, and Tumstatin, derived from the non-collagenous domain of the α3 chain of type IV collagen. Both molecules are generated from vascular basement membrane components and inhibit pathological angiogenesis and tumor growth in experimental models^5-7^.

Although originally grouped together as extracellular matrix-derived angiogenesis inhibitors, Endostatin and Tumstatin were reported to regulate endothelial cell function through distinct mechanisms. We previously demonstrated that Tumstatin functions as an endothelial cell-specific inhibitor of protein synthesis by binding αVβ3 integrin and suppressing FAK, PI3K, Akt, and mTOR signaling, resulting in inhibition of cap-dependent translation and induction of endothelial cell apoptosis^8^. Subsequently, we reported that human Tumstatin and human Endostatin exhibit fundamentally different antiangiogenic activities: Tumstatin inhibits endothelial proliferation and promotes apoptosis through αVβ3 integrin-dependent suppression of the PI3K/Akt/mTOR/4E-BP1 pathway, whereas Endostatin inhibits endothelial migration through α5β1 integrin-dependent regulation of MAP kinase signaling with little or no effect on Akt activation^9^. Subsequent investigations further supported the biological importance of the Tumstatin-Akt pathway and collectively shown that Tumstatin-derived fragments inhibit endothelial proliferation, induce apoptosis, suppress neovascularization, and reduce tumor progression in multiple experimental systems^5,6^. Here, we independently evaluated the differential effects of recombinant human Tumstatin and recombinant human Endostatin on Akt phosphorylation in human endothelial cells using Human umbilical vein endothelial cells (HUVEC) grown in in vitro on fibronectin substrate.

## Materials and Methods

Human umbilical vein endothelial cells (HUVEC; Lonza, C2519A, Batch 25TL191608) were cultured in complete EGM-2 medium (Lonza) at 37°C in a humidified atmosphere containing 5% CO_2_. Cells between passages 2 and 4 were used for all experiments. HUVEC were plated onto fibronectin-coated tissue culture dishes (Corning BioCoat Fibronectin, #354402) and incubated in serum free medium in the presence or absence of recombinant human Tumstatin or recombinant human Endostatin (20 μg/mL) for the indicated time intervals. Recombinant human Tumstatin (rh-Tum) and recombinant human Endostatin (rh-Endo) were generated as previously described^8,9^. Cell lysates were collected and analyzed by western blotting using antibodies against phospho-Akt (Cell Signaling Technology #4060, 1:200 dilution) and total Akt (Cell Signaling Technology #9272; 1:200 dilution). Horseradish peroxidase-conjugated secondary antibodies were used for signal detection. Ladder: Biorad Precision Plus (1610374).

## Results

### Differential Regulation of Akt Phosphorylation by Tumstatin and Endostatin

To determine whether Tumstatin and Endostatin differentially regulate Akt signaling in endothelial cells, serum-starved HUVEC were plated on fibronectin-coated dishes and exposed to recombinant proteins under conditions analogous to those previously reported^9^. Western blot analysis demonstrated that recombinant human Endostatin did not substantially alter Akt phosphorylation relative to untreated controls at the evaluated experimental timepoints (**Figure 1**). In contrast, exposure to recombinant human Tumstatin resulted in reduced phospho-Akt levels while total Akt expression remained unchanged (**Figure 1**). The selective decrease in phospho-Akt observed in Tumstatin-treated cells indicates inhibition of Akt activation rather than reduced Akt abundance. These findings closely mirror the signaling pattern previously reported in which Tumstatin inhibited sustained Akt phosphorylation whereas Endostatin failed to suppress Akt activation in endothelial cells cultured on fibronectin^9^. Together, these data provide independent confirmation of a central mechanistic distinction between these two collagen-derived angiogenesis inhibitors.

**Figure 1.**
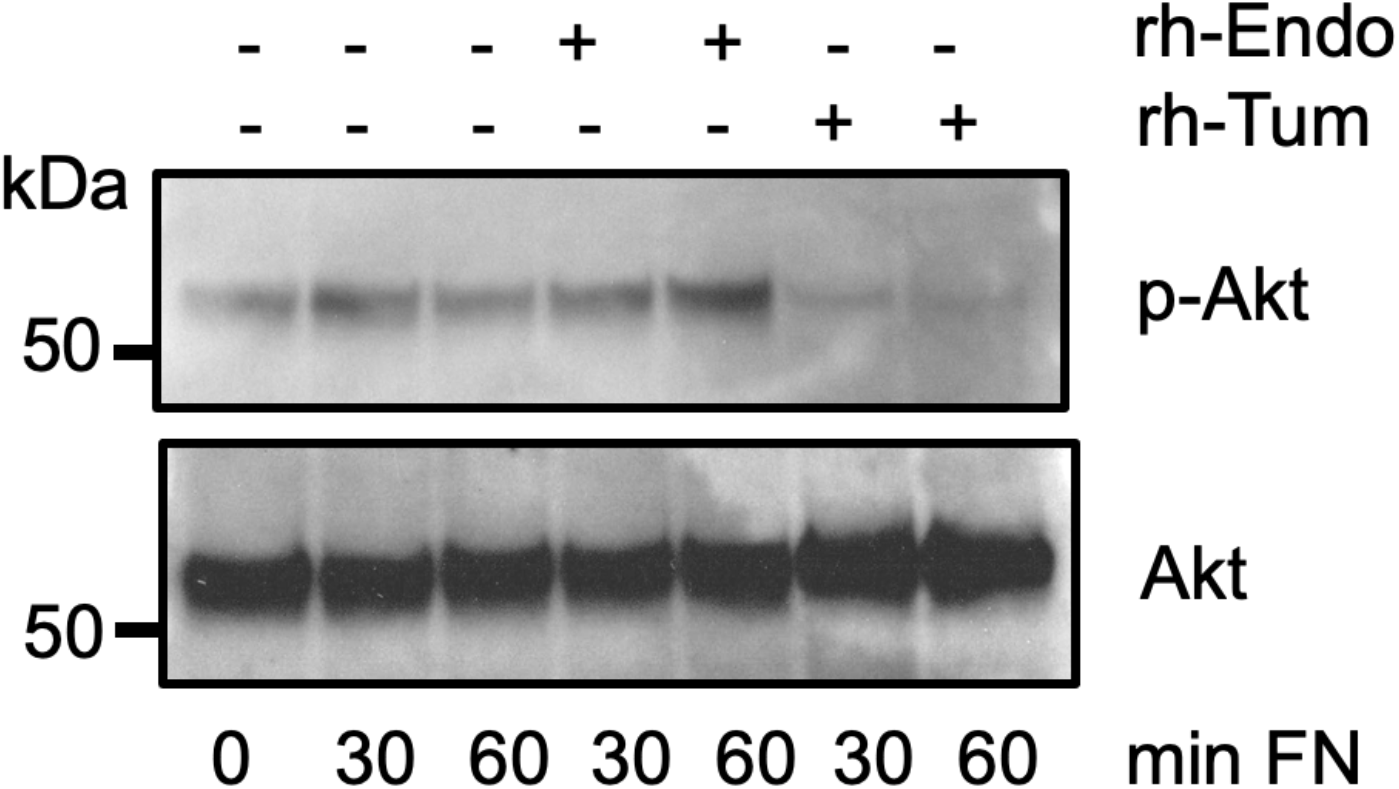
Effect of rh-Tum and rh-Endo on AKT signaling. Serum-starved HUVEC were plated on Fibronectin (FN)-coated dishes and treated with rh-Endo (20μg/mL) or rh-Tum (20μg/mL) for the indicated times (min: minutes) and lysates analyzed by Western blot. Immunoblots of phosphorylated Akt (p-Akt, upper) and total Akt (lower) demonstrate no changes in phosphorylation on rh-Endo treatment, whereas rh-Tum significantly inhibited the phosphorylation of Akt. rh-Endo: recombinant human Endostatin; rh-Tum: recombinant human Tumstatin; kDa: kilodalton.

## Discussion

The present study independently validates a key mechanistic observation in the angiogenesis field. We report here that Tumstatin suppresses endothelial Akt signaling in HUVEC whereas Endostatin does not, similar to previous published studies^8,9^. This distinction remains biologically important because Akt functions as a central regulator of endothelial survival, protein synthesis, proliferation, and angiogenic responses. Previous studies established that Tumstatin engagement of αVβ3 integrin suppresses the FAK/PI3K/Akt/mTOR signaling cascade, resulting in inhibition of cap-dependent translation and endothelial apoptosis^8,9^. In contrast, Endostatin inhibits endothelial migration through α5β1 integrin-mediated signaling involving FAK, c-Raf, MEK, ERK1, and p38 MAP kinase pathways while leaving Akt activation largely intact^9^. The preservation of Akt phosphorylation observed in the present study therefore represents an expected mechanistic pathway of Endostatin signaling. Our results are consistent with later studies demonstrating the broader significance of Akt suppression in Tumstatin biology, specifically that inhibition of the PTEN/Akt pathway in glioma cells contributes directly to αVβ3-dependent growth suppression and antiangiogenic activity mediated by Tumstatin-derived peptides^10^.

Additional investigations have demonstrated that Tumstatin-derived fragments inhibit endothelial proliferation, induce apoptosis, and suppress tumor-associated angiogenesis in diverse experimental models^5,6^. Our results reinforce the concept that endogenous extracellular matrix fragments regulate angiogenesis through distinct integrin-specific signaling programs rather than through a common antiangiogenic mechanism.

## Acknowledgments

We are grateful to M. Kirtley and A. Comptdaer for assistance in procuring reagents and reviewing methodology.

## Funding

This work was supported by Kalluri laboratory funds available to RK.

## Conflict of Interests

The authors declare no conflict of interest.

## Notes

### Competing Interest Statement

The authors have declared no competing interest.

